# Single-cell integrative analysis of CAR-T cell activation reveals a predominantly T_H_1/T_H_2 mixed response independent of differentiation

**DOI:** 10.1101/435693

**Authors:** Iva Xhangolli, Burak Dura, GeeHee Lee, Dongjoo Kim, Yang Xiao, Rong Fan

## Abstract

We present the first comprehensive portrait of single-cell level transcriptional and cytokine signatures of anti-CD19 4-1BB/CD28/CD3ζ CAR-T cells upon antigen-specific stimulation. Both CD4^+^ ‘helper’ and CD8^+^ cytotoxic CAR-T cells are equally effective in directly killing target tumor cells and their cytotoxic activity is associated with the elevation of a range of T_H_1 and T_H_2 signature cytokines (e.g., IFNγ, TNFα, IL5, and IL13), as confirmed by the expression of master transcription factors TBX21 (T-bet) and GATA3. However, rather than conforming to stringent T_H_1 or T_H_2 subtypes, single-cell analysis reveals that the predominant response is a highly mixed T_H_1/T_H_2 function in the same cell and the regulatory T cell (T_reg_) activity, although observed in a small fraction of activated cells, emerges from this hybrid T_H_1/T_H_2 population. GM-CSF is produced from the majority of cells regardless of the polarization states, further contrasting CAR-T to classic T cells. Surprisingly, the cytokine response is minimally associated with differentiation status although all major differentiation subsets such as naïve, central memory, effector memory and effector are detected. All these suggest that the activation of CAR-engineered T cells is a canonical process that leads to a highly mixed response combining both type 1 and type 2 cytokines together with GMCSF, supporting the notion that ‘polyfunctional’ CAR-T cells correlate with objective response of patients in clinical trials. This work provides new insights to the mechanism of CAR activation and implies the necessity for cellular function assays to characterize the quality of CAR-T infusion products and monitor therapeutic responses in patients.

## INTRODUCTION

Adoptive transfer of anti-CD19 Chimeric Antigen Receptor (CAR)-T cells has demonstrated remarkable efficacy in treating patients with B cell acute lymphoblastic leukemia (B-ALL), chronic lymphocytic leukemia (CLL) and other indolent lymphomas^1–4^. Despite the demonstrated success, there exist large variation of responses and unpredictable toxicity in patients^5, 6^, which could be attributed in part to inter-patient and intra-population heterogeneity of CAR-T infusion product, requiring a higher resolution approach to characterize not only phenotypic composition but the function of CAR-T cells at the systems level. The scFv ectodomain of CAR binds to CD19 expressed on the surface of a target tumor cell and transmits signal via the transmembrane linker to the intracellular signaling domain such as CD3ζ to elicit T-cell activation independent of signaling from T-cell receptor (TCR)-mediated binding to peptide major histocompatibility complex (p-MHC)^7^. Incorporating a co-stimulator domain such as CD28 or 4-1BB (CD137) in the second generation CAR further enhanced proliferation, persistence, and potency^8^. Therefore, the mechanism of CAR-T activation could differ substantially from that of classic T cells. But it remains inadequately characterized as of today. The questions like how different CAR-T cell subsets such as CD4 helper vs CD8 cytotoxic cells respond to CAR stimulation, how the polarization subtypes such as T_H_1, T_H_2, T_reg_, etc. differentially control CAR-T cell responses, how the differentiation status could affect the activation state, are all yet to be fully elucidated.

Current methods for evaluating CAR-T cell activation include the measurement of IFNγ secretion by ELISA or the detection of IFNγ-secreting cells by ELISpot^9, 10^. Multiparameter flow cytometry was used for immunophenotyping of CAR-T cells, which is one of the mainstay techniques used for monitoring CAR-T product manufacturing, but the number of markers and functions (cytokines) it can measure is limited^11, 12^. Intra-cellular cytokine staining for flow cytometric analysis is not a true secretion assay and often leads to over-estimation of cytokine-secreting cells. Recently, Xue *et al.* used single-cell multiplex cytokine profiling to measure the cytokine output of CD19-BB-3z CAR-T cells using CD19-coated beads and revealed a diverse polyfunctional response upon activation^13^. However, the cytokine profile of a CAR-T cell is yet to be directly correlated to cytotoxicity, subtype and signaling in order to elucidate the underlying mechanisms.

Herein, we use high-throughput single-cell 3’ mRNA transcriptome sequencing, single-cell multiplex cytokine secretion assay, together with live cell imaging of cytotoxic activity to interrogate third-generation anti-CD19 4-1BB/CD28/CD3ζ (CD19-BB-28-3z) CAR-T cells, yielding the first comprehensive portrait of single-cell level transcriptional and cytokine signatures of CAR-T cells upon antigen-specific stimulation. A range of cytokines are produced in single CAR-T cells in direct correlation with cytotoxicity, irrespective of CD4^+^ ‘helper’ vs CD8^+^ cytotoxic cell types. Single-cell analysis reveals that the predominant response is a highly mixed T_H_1/T_H_2 function in the same cell that also produce granulocyte-macrophage colony stimulating factor (GM-CSF). Regulatory T cell (T_reg_) activity, although observed in a small fraction of activated cells, emerges from this hybrid T_H_1/T_H_2 population. Surprisingly, the cytokine response is minimally associated with differentiation status although all major subsets such as naïve, central memory, effector memory, and effector cells are detected. Our study suggests that the activation of CAR-engineered T cells is a canonical process associated with a highly mixed multi-functional response, supporting the notion that ‘polyfunctional’ CAR-T cells correlate with objective response of Non-Hodgkins Lymphoma patients reported in a CD19 CAR-T clinical trial^14^.

## RESULTS AND DISCUSSIONS

### Study design: single-cell level measurement of transcriptional, cytokine, and cytotoxic function

We combined a set of single-cell techniques to interrogate CAR-T cells upon antigen specific stimulation (**Figure 1a**) in order to quantitatively dissect the correlation of activation states with T-cell subtypes, differentiation, and other factors such as intracellular signaling cascades, co-stimulators and immune checkpoints. CAR-T cells used in this study were manufactured through *ex vivo* transduction of autologous T cells with a CD19-BB-28-3z CAR construct, expansion for ~10 days using CD3/CD28 Dynabeads, and then purification by bead removal and enrichment for CAR expression. T cells were isolated from three healthy donors and for the purpose of this study the human B-cell lymphoma Raji cell line was used as a target. Single-cell 3’ mRNA transcriptome profiling was performed using a massively parallel cell barcoding method^15^ implemented in a bead-in-a-well microchip. Single CAR-T cell cytolytic activity was measured by co-seeding CAR-T cells and target tumor cells in the microwell array to image the uptake of SYTOX Green nucleic acid dye^16^ indicative of target cell lysis by CAR-T cells. Single-cell multiplex cytokine secretion assay was performed using a previously developed antibody barcode microchip assay^17–19^, which was further modified in this study to co-measure cytotoxicity by seeding both CAR-T and tumor cells in the same single-cell protein secretion assay microchamber. Integrating all these single-cell analyses tools provides an unprecedented resolution to correlate the activation states such as anti-tumor cytotoxicity and cytokine response in single CAR-T cells to their phenotype, differentiation, signaling, and other characteristics.

**Figure 1.**
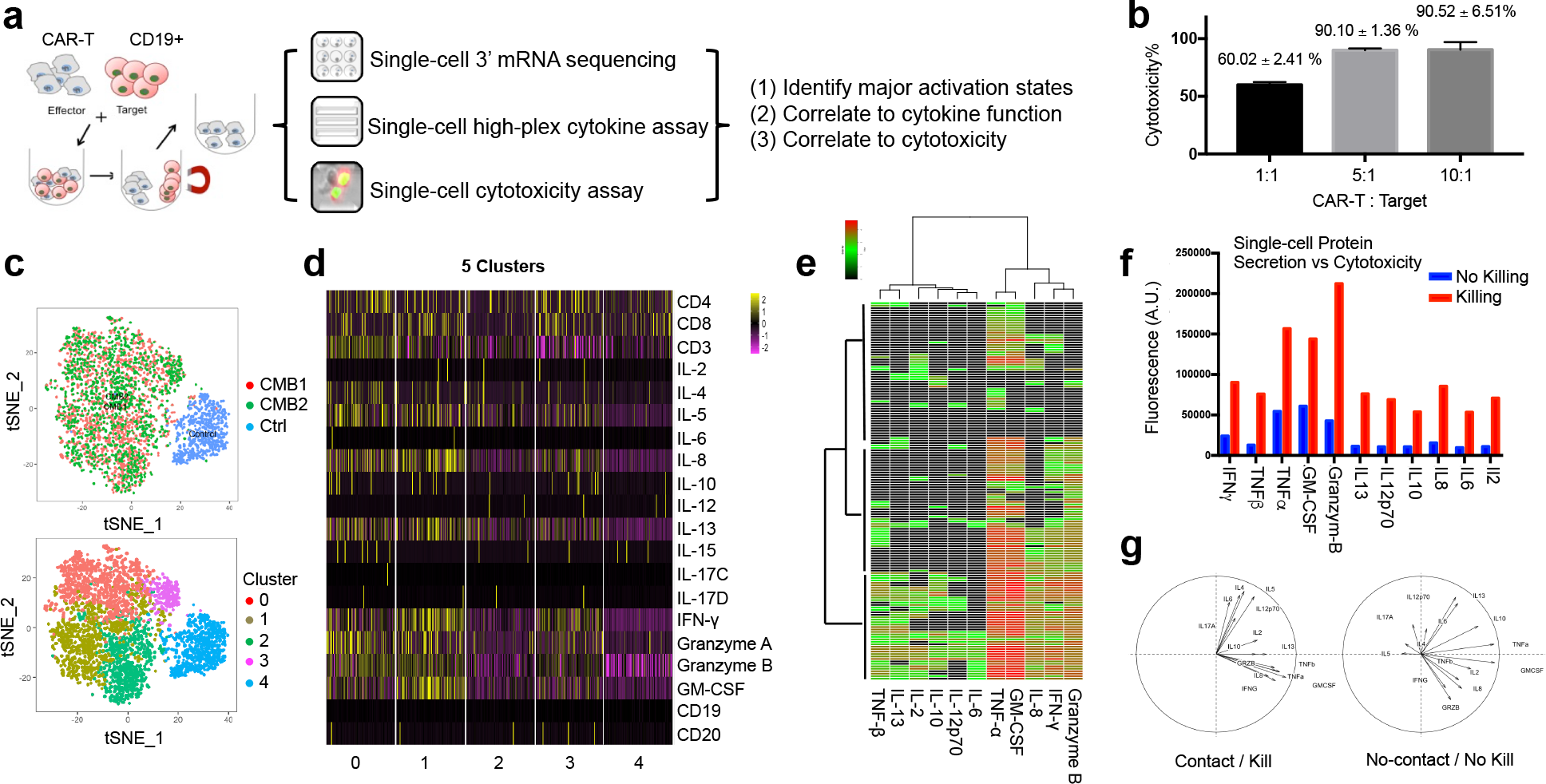
Single-cell integrative analysis reveals varying degrees of CAR-T cell activation and the direct correlation to cytotoxicity. (a) Approach – single-cell transcriptome, cytokine secretion, and live cell tracking of single-cell cytolystic activity to characterize CAR-T cells. (b) LDH assay confirms both CD4 and CD8 CAR-T cells are cytotoxic to CD19-expressing tumor cells. (c) tSNE analysis of single-cell transcriptomes from a duplicate activation experiment (CMB1&CMB2) vs unstimulated control (Ctrl), showing highly consistent and efficient activation of >95% CAR-T cells and four major/clusters states (0-3) with varying degrees of activation compared to control (5). (d) Major cytokine gene expression in single cells from major clusters (0-4). (e) Single-cell multiplex protein secretion assay reveals three ties of activity, each of which is a different combination of cytokines shown by hierarchical clustering using Manhattan distance. Each row represents a microwell with a successful target cell killing event and each column corresponds to a protein of interest. Color intensity correlates with log scale of normalized fluorescence intensity of cytokine detected. (f) Cytokine secretion intensity is elevated in microchambers where a CAR-T cell killed target tumor cells. [y-axis: Background normalized fluorescence intensity of detected cytokines] (g) PCA shows distinct groups of cytokine secretions between killing and no-killing cases.

### CAR activation leads to heterogeneous responses involving a range of cytokines, correlating directly to cytotoxicity

Prior to conducting single-cell analyses, a population LDH assay demonstrated these CAR-T cells are highly cytotoxic (60.02% at 1:1 and 90.1% at a 5:1 CAR-T:Target ratio) (**Figure 1b**), confirmed by time-lapse imaging of SYTOX Green uptake (**Supplementary Figure 1a**), and concurrently correlated with the secretion of a panel of cytokines (*e.g.*, IFNγ, TNFα, GMCSF, IL4, IL5, IL8, and IL13) detected by a protein microarray assay (**Supplementary Figure 1b**). Single-cell massively parallel 3’ mRNA sequencing was conducted with unstimulated alone CAR-T cells (Ctrl) and stimulated CAR-T cells (replicates: CMB1 and CMB2). The stimulated cells were prepared by co-culturing with CD19-expressing target cells for 6 hrs and then purified by magnetically cell sorting to remove target cells). The sequencing data were of high quality (**Supplementary Figure 2&3**) and 3817 single CAR-T cell transcriptomes were obtained, which were visualized with t-Distributed Stochastic Neighbor Embedding (t-SNE) plots (**Figure 1c upper panel**). We observed high consistency between replicates and highly efficient activation with <0.5% of activated CAR-T cells (red and green) residing in the Ctrl cluster (blue). Differential gene expression analysis (**Supplementary Figure 4 and Figure 1c low panel**) further identified four clusters 0-3 within the activated population that show varying degrees of cytokine responses (**Figure 1d**). Most cytokines measured are highly expressed in clusters 0 and 1, moderately expressed in cluster 2, and very sparsely in cluster 3 that is barely above control (cluster 4). Single-cell 14-plex cytokine secretion assay confirms the heterogeneous activation with 3 clusters identified that secret most, nearly half, or very few cytokines, respectively (**Figure 1e**). To correlate cytokine function directly to cytotoxicity, live cell imaging of SYTOX Green uptake was performed in the CAR-T / Raji cell pairs co-loaded in single-cell cytokine secretion microchambers over 12 hours, and the cytokine profiles are separated into no-killing vs killing groups (**Supplementary Figure 5 and Figure 1f**). Rather than specific cytotoxic effector cytokines such as Granzymes, a range of cytokines are elevated upon CAR activation and directly correlate to cytotoxic activity. Finally, principal component analysis (PCA) revealed the group of cytokines into two clusters, one dictated by TNFα, IFNγ, Granzyme B, GMCSF and IL8, and the other dictated by IL4, IL5, IL6, and IL17. In contrast, the cytokine profiles from single CAR-T cells that did not kill tumor cells showed no distinguishable grouping among these cytokines (**Figure 1g**). A diverse landscape of antigen-specific response was observed in CD19-BB-3z CAR-T cells^13^. Their dominant polyfunctional cells also produce GranzymeB, GM-CSF, IL8, TNFα, IL13, and IFNγ, in concordance with our observation in the third-generation CAR-T cells.

### Both CD4^+^ and CD8^+^ CAR-T cells are highly cytotoxic and produce similar combination of cytokines at the proteomic and transcriptional level

In addition to the CD4 differentiation subsets discussed above, cytotoxic CD4^+^ T cells (CTL CD4), able to secrete Granzyme B and perforin, have been observed during virus infections and antitumor and chronic inflammatory responses^20^. The percentage of CTL CD4 cells within the whole CD4^+^ T-cell population is very small, but in the case of a mixed CD4+ and CD8+ CAR T-cell population, 66 % of all cells secrete Granzyme B. To examine whether the varying degrees of CAR-T cell activation is a consequence of varying CD4 helper cell phenotype present in the CAR-T product, we conducted SYTOX Green assay with target cells loaded in microfabricated wells (100μm×100μm) together with CD4^+^ or CD8^+^ CAR-T cells, respectively (**Figure 2a**and **Supplementary Movies**). Both CD4+ and CD8+ CAR-T cells are equally efficient in killing target cells^21^. The time and distance for a CD4^+^ or CD8^+^ CAR-T cell to find and lyse target tumor cells are similar. Single-cell cytokine secretion (proteome) and mRNA expression (transcriptome) data showed nearly indistinguishable frequency and level of cytokine expression between CD4^+^ and CD8^+^ subsets (**Figure 2b**) except a slightly increased level of Granzyme B in CD8^+^ cells and a slightly upregulated expression of IL4, IL5, IL13, and IL10 in CD4+ cells. Both subsets are highly polyfunctional (>50% co-producing 5+ cytokines and the polyfunctionality in CD4^+^ subset is slightly higher). PCA further confirms a similar pattern of cytokine grouping in both subsets. This is consistent with previous findings^13^ and supports the hypothesis that CAR activation is largely independent of class 1 vs 2 p-MHC signaling. CAR-T cells with defined CD4:CD8 composition have been investigated to treat B-ALL patients and demonstrated high potency and allowed for delineating factors correlated with expansion, persistence and toxicity^22^. Our results confirmed the importance of CD4 cells in CAR-T product and the necessity to fully evaluate the role of these “helper” T cells in immunotherapy.

**Figure 2.**
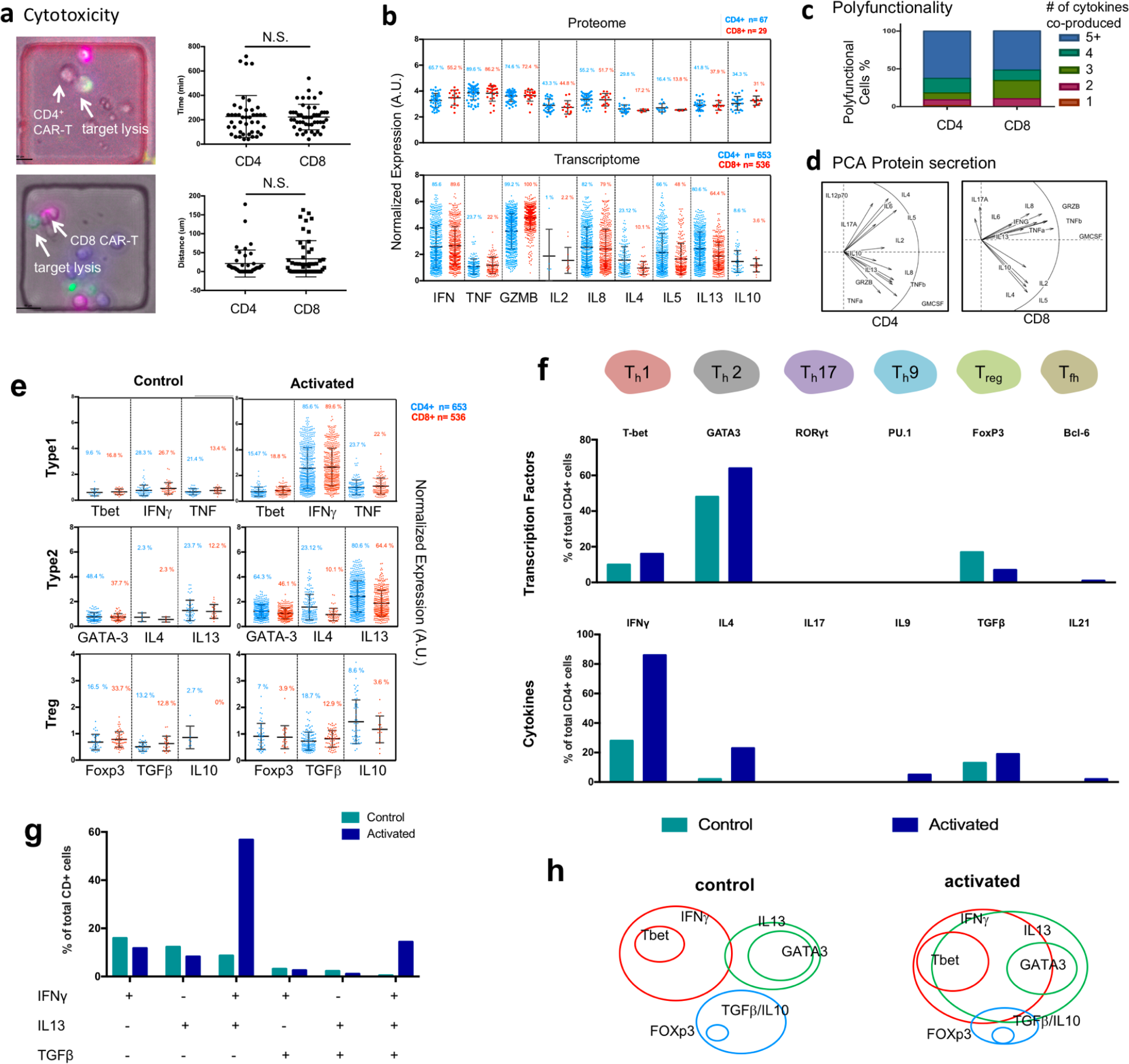
Mapping single cell data to phenotypes reveals a highly mixed T_H_1/T_H_2 cell response. (a) Single-cell cytolystic activity assay reveals that both CD4 and CD8 CAR-T cells are cytotoxic and show insignificant difference in anti-tumor reaction. (b) Comparing cytokine gene expression (transcriptome) and secretion (proteome) in single CAR-T cells between CD4 and CD8 subsets. (c) Polyfunctionality, which defines the abiliyt for a single cell to co-secrete multiple cytokines, is slightly higher in CD4 relative to CD8 CAR-T cells. (d) PCA reveals similar clustering of major cytokine secretions between CD4 and CD8 cells. (e) Expression type 1, type 2, and regulatory T cell signature genes in single CAR-T cells upon activation. (f) Quantification of CD4 T cell subsets according to signature transcription factor and cytokine gene. (g) Analysis of co-expression of Th1, Th2, and Treg cytokines in single CAR-T cells. (h) Diagram showing phenotypic composition of CD4+ CAR-T cells and the alteration upon antigen-specific activation.

### Mapping into subtypes reveals a predominantly mixed T_H_1/T_H_2 response in conjunction with T_reg_ activity in the same CAR-T cells

To access whether the observed heterogeneous activation depends on CAR-T cell polarization and subtypes, we examined the expression of master transcription factors(TFs) and signature cytokines. Both type 1 and 2 TFs are elevated and their respective signature cytokines produced (**Figure 2e**). Foxp3, a master TF for regulatory T cells, is present in unstimulated cells, downregulated upon activation, and associated with elevated production of regulatory cytokines TGFβ and IL-10. All these are minimally dependent on CD4 vs CD8 despite a slightly higher Type 2 activity in CD4^+^ cells. Further mapping all single CD4 cell transcriptomes into T_H_1, T_H_2, T_H_17, T_H_9, T_reg_, and T_FH_ subtypes showed that T_H_1 and T_H_2 are dominant and the T_reg_ response is observed (**Figure 2f**). Although T-bet is detected in 15.5% of cells, IFNγ is produced from 85.6% of cells, suggesting the interferon response is also elicited or amplified by T-bet-independent pathways, for example, STAT1. GATA3 is expressed in 64.3% of cells but the predominant T_H_2 cytokine IL4 is produced by 23.1% of these cells whereas other T_H_2 cytokines (66% IL5+ and 80.6% IL13+) are more predominant, indicative a prevalent but slightly skewed T_H_2 response, consistent with previous report that IL13 rather than IL4 is the dominant T_H_2 cytokine observed in CD19-BB-3z CAR-T cell activation^13^. Thus, for further analysis IL13 rather than IL4 was chosen as the T_H_2 dominant cytokine.

To further answer if these are stringent polarization subtypes as in classic T cell biology, we quantified the frequency of dominant T_H_1 (IFNγ), T_H_2(IL13), and T_reg_ (TGFβ) responses in the same single cells. It turned out the majority of CD4^+^ CAR-T cells upon activation show a mixed T_H_1 (IFNγ) and T_H_2 (IL13) response and the majority (75%) of T_reg_ (TGFβ) - like cells are triple positive for T_H_1 (IFNγ) and T_H_2(IL13) **(Figure 2g)**. Therefore, CAR-T cell activation appears to differ substantially from classic T cells in that the predominant response is a highly mixed T_H_1 / T_H_2 phenotype, within which a fraction of cells further exhibit regulatory activity, presumably to counteract over-activation (**Figure 2h**). Hegazy *et al.* found that interferons can direct T_H_2 cell reprogramming to generate a hybrid subset with combined T_H_2 and T_H_1 cell functions^23^, which could be a mechanism in our CAR-T cells given that GATA3 is prevalently expressed and interferon response is the most profound, which together may confer T_H_1 function to GATA3^+^ CAR-T cells. Other mechanisms reported include stochastic cytokine expression leading to the formation of mixed T helper cell states^24^. Peine *et al.* reported that stable T_H_1/T_H_2 hybrid cells arise *in vivo* from naïve precursors to limit immunopathologic inflammation^25^, which may explain in part the notion that CAR-T cells in general show a balanced response in anti-tumor effector and inflammatory toxicity despite the unusually wide range of cytokines co-produced per cell.

### GM-CSF is highly prevalent in activated CAR-T cells and co-produced with Type 1 and 2 cytokines

GM-CSF is a potent proinflammatory cytokine involved in the recruitment, maturation and activation of myeloid cells^26^. The helper T cells producing GM-CSF have been identified and recently reported to serve a nonredundant function in autoimmune pathogenesis, arguably representing a unique helper T cell subset^27^. Our data showed that >80% of activated CD4^+^ cells express GM-CSF, confirmed independently by single-cell cytokine secretion (proteome) and single-cell mRNA sequencing (transcriptome) (**Supplementary Figure 6**). GM-CSF production is associated with both Type 1 (IFNγ) and Type 2 (IL5) responses. We observed that 89% of T-bet^+^ cells and 83.5% of GATA3^+^ cells produce GM-CSF. Previously, it was found that STAT5 programs a distinct subset of GM-CSF-producing T_H_ cells in neuroinflammation and autoimmunity^28^. In this work, 65.2% of GM-CSF-expressing CD4^+^ CAR-T cells are negative or low for STAT5. Instead, we observed the expression of a wide range of STATs, but STAT1 is predominant, which may contribute to the profound T_H_1 interferon response despite a modest level of T-bet expression. However, Gene Set Enrichment Analysis (GSEA) confirmed the relevance of STAT5 signaling and also revealed the importance of IL6/STAT3 (**Supplementary Figure 7**), which has been observed as a determinant of CAR-T memory phenotype and therapeutic persistence in patients^29^. Finally, the JAK/STAT signaling relies on phosphorylation of STATs and their nuclear translocation. The mechanism leading to a prevalent GM-CSF-producing phenotype in activated CAR-T cells is yet to be further investigated.

### Cytokine production in CAR-T cells upon antigen-specific activation is minimally dependent on differentiation status

To examine if the activation state of CAR-T cells is dependent on the differentiation status, we stratify the transcriptomes of single cells (both control and activated) into four subsets: naïve (T_N_), central memory (T_CM_), effector memory (T_EM_), and effector (T_EFF_), based on the markers CD45RA and CCR7. The naïve T_N_ subset herein may also contain T stem-cell memory (T_SCM_) cells. We found that both CD4^+^ and CD8^+^ populations are mainly comprised of naïve and effector cells but central memory and effector memory cells were also observed (**Figure 3a**). CAR activation leads to increased frequencies of T_CM_ and T_EM_ cells and as a result a reduced frequency of T_N_ cells. However, correlating the expression of a range of stimulatory and cytotoxic cytokines in CD4^+^ cells involved in T_H_1, T_H_2, T_reg_ phenotypes to differentiation status demonstrates none or minimal dependence (**Figure 3b**). A closer look revealed slightly increased TNFα expression and decreased T_H_2 cytokines (e.g., IL4 and IL13) in the T_CM_ subset (red bars in **Figure 3b**). CAR-T product containing central memory cells has been reported as correlating with *in vivo* expansion, persistence and potency^30^. Thus, the role of T_CM_ subset in CAR-T activation requires further investigation. T-cell growth factor receptors IL2RA (CD25) and IL15R show no statistically significant difference, either. Interestingly, CD4^+^CD25^+^ T cells are considered as immunosuppressive and often used as a surrogate for Foxp3^+^ T_reg_ cells^31^. In this study, nearly 100% of CAR-T cells upon activation are CD25-positive but apparently do not exert the suppressor function, highlighting the importance to characterize the function of CAR-T cells following activation in addition to surface marker phenotyping. It is worth noting that the differentiation status of CAR-T cells manufactured from healthy donors and patients with B-cell malignancies could differ substantially and the patient samples can contain significantly higher percentage of central and effector memory cell subsets^22^. However, our work can still teach the fundamental mechanism of CAR-T cell response as a function of differentiation status as we can identify all the major differentiation subsets (*e.g.*, T_N_, T_CM_, T_EM_, and T_EFF_).

**Figure 3.**
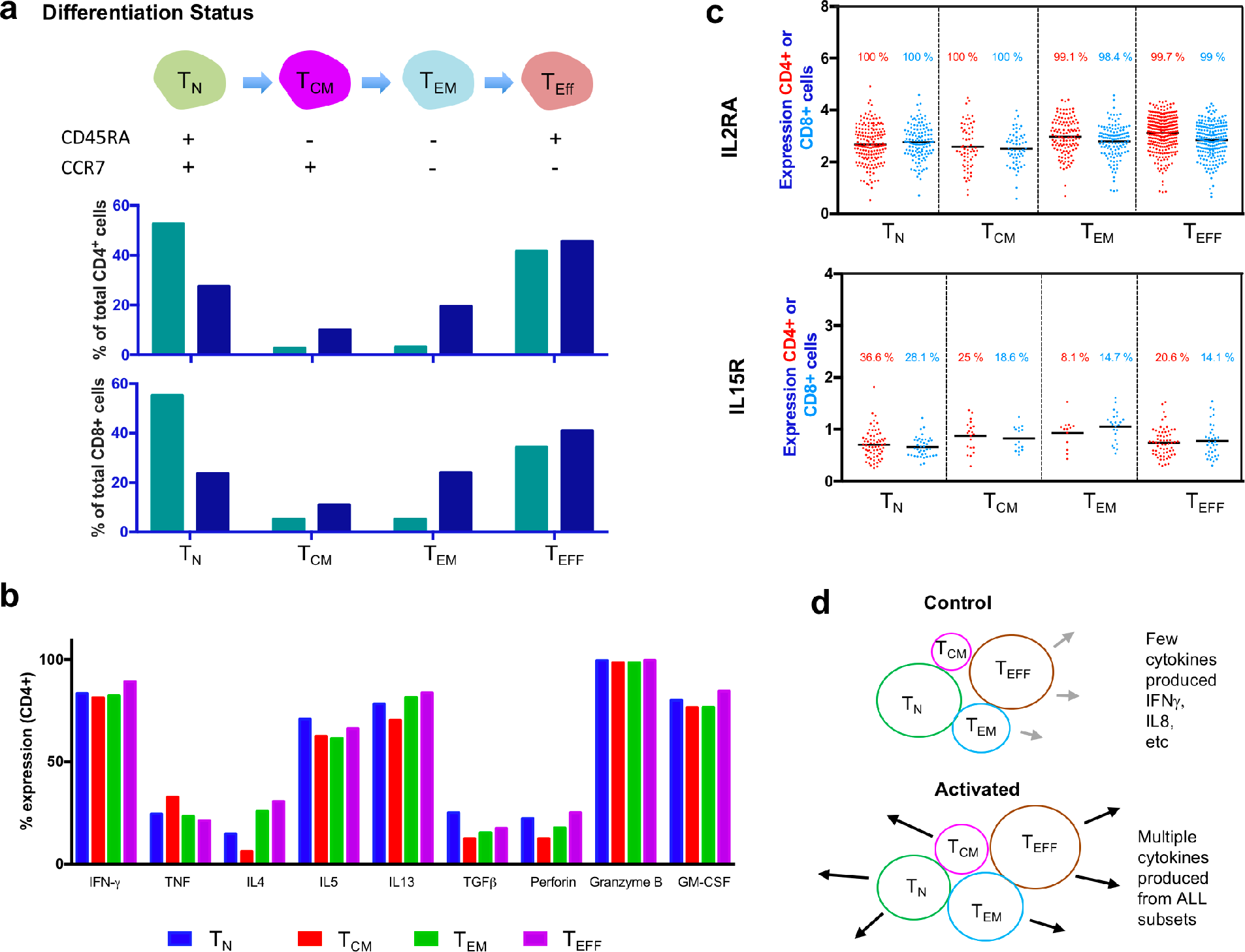
Correlating differentiation status of CAR-T cells to cytokine functions. (a) Stratification of unstimulated and stimulated CAR-T cells into different differentiation status using a pair of markers CD45RA and CCR7. (b) Correlating the expression of a range of cytokines in CD4^+^ cells to differentiation status. (c) Expression of T-cell growth factor receptors IL2RA and IL15R in different subsets of CAR-T cells upon activation. (d) A model to describe the activation of CAR-T cells to produce effector cytokines is largely independent of differentiation status.

### CAR activation leads to elevated expression of CTLA4 that correlates to upregulation of IL10 in CD4^+^ and TGFβ in CD8^+^ subsets

To investigate the possible relationship between CAR-T cell activation and the changes in co-stimulators and immune checkpoints, we examined their expression in single-cell transcriptomes. We observed that CTLA4 is the most upregulated among all immune checkpoints upon CAR activation (**Figure 4a**). PD-1 demonstrates a similar trend but its expression level is low and the frequency is only 9.8% in CD4^+^ and 3.7% in CD8^+^ subsets. PD-1 is also the primary marker of T cell exhaustion^32^ and thus our data suggest an infrequent incidence of acute stimulation-induced exhaustion following CAR-T cell action. Concurrently there is a decreased frequency of ICOS and OX40 expression but slightly increased mRNA levels in a fraction of cells. Correlative analysis revealed that unstimulated cells are predominantly ICOS^+^CTLA4^-^ but the activated cells contain all three combinations and the ICOS+CTLA4+ subset is dominant in CD4^+^ CAR-T cells(**Figures 4b&c**). Finally, the question is whether the significant upregulation of CTLA4 correlates or may contribute to the observed cytokine response heterogeneity. Comparing CTLA4^high^ (top 10%) cells to CTLA4^low^ (bottom 10%) cells for a range of cytokines (**Supplementary Figure 8**), we observed significant increase of IL10 expression in CD4^+^ subset and TGFβ expression in CD8^+^ subset whereas all other cytokines are minimally or insignificantly correlated with CTLA4, suggesting a possible mechanism for CAR-T cells to control immune homeostasis without developing distinct regulatory subtype, which has important implication in combining checkpoint blockage and CAR targeted immunotherapies^33–35^.

**Figure 4.**
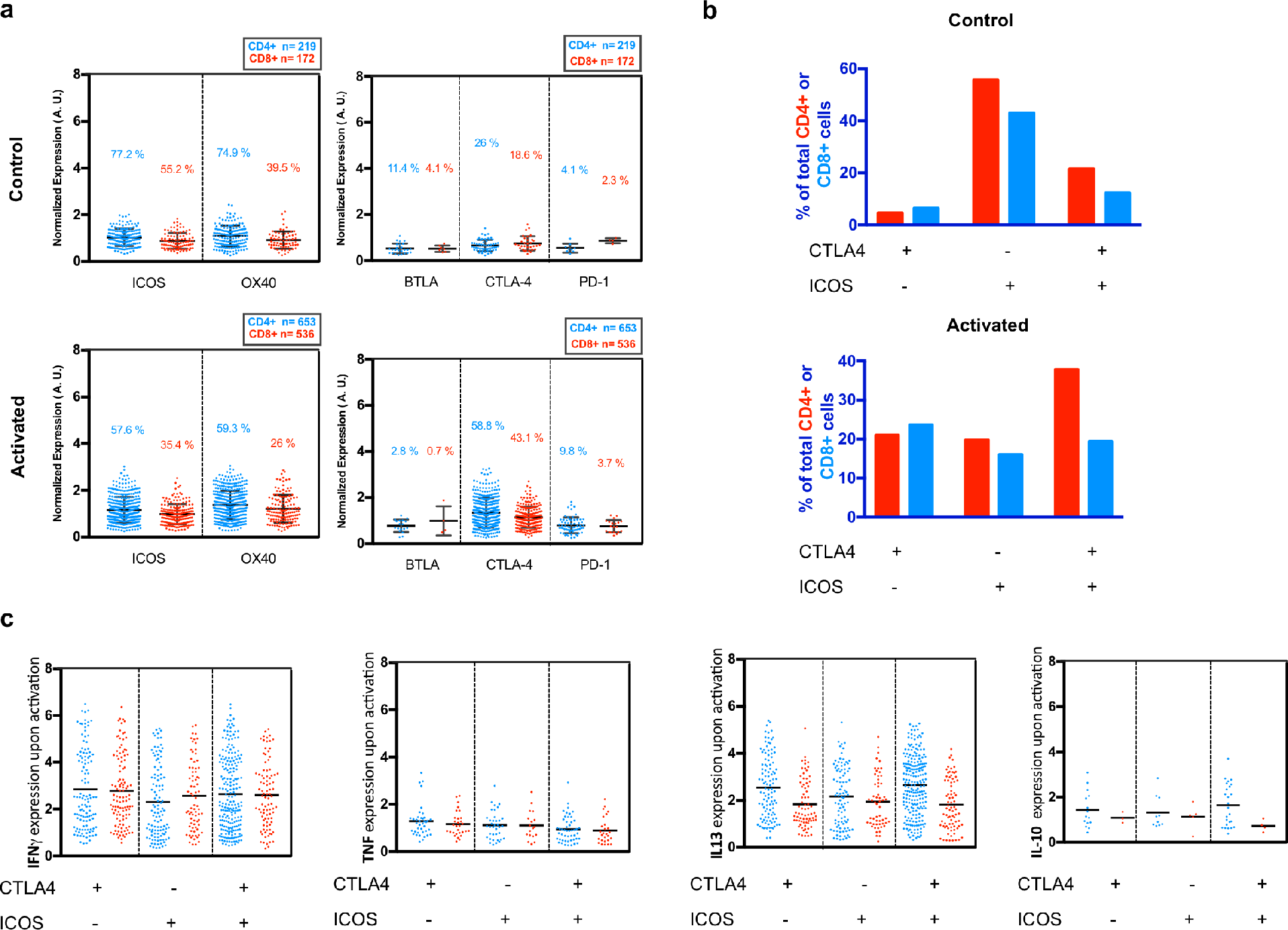
Effect of activation state on co-inhibitory (BTLA, CTLA-4, and PD-1) and co-stimulatory receptors (ICOS and OX40). (a) Common co-stimulatory and co-inhibitory expression profiles in CD4 and CD8 subpopulations of activated and control CAR-T cells. In the CD4+ subset population, CTLA-4 expression doubles from the control (26%) to the activated state (58.8%) while there is slight decrease in the co-stimulatory ICOS expression from 77.2 % in the control state to 57.6% in the activated state. (b) Percentage of cells expressing three combinations of CTLA-4 and ICOS are plotted. (c) Each cell population from b was separated and the expression of Th1 (IFNγ, TNFα), Th2 (IL13), and Treg (IL10) was plotted.

## CONCLUSIONS

We use high-throughput single-cell 3’ mRNA transcriptome sequencing, multiplexed single-cell cytokine secretion assay, together with live cell imaging of cytolytic activity to interrogate third-generation anti-CD19 4-1BB/CD28/CD3ζ (CD19-BB-28-3z) CAR-T cells upon antigen-specific activation. CD4^+^ and CD8^+^ CAR-T cells are found to be equally effective in direct killing of target tumor cells and the cytotoxic activity is associated with the elevated co-production of a wide range of cytokines. Both T_h_1 and T_h_2 responses are prevalent, as confirmed by master transcription factors T-bet and GATA3 as well as signature cytokines IFNγ, TNFα, IL5, and IL13. Regulatory T cell activity, although detected in a small fraction of cells, is associated with elevated TGFβ and IL10 production. But unexpectedly, all these responses are often observed in the same CAR-T cells rather than distinct subsets, supporting the notion that polyfunctional CAR-T cells correlate with objective response of patients in clinical trials(14). GM-CSF is produced from the majority of cells regardless of the polarization state, further contrasting CAR-T to conventional T cells. We found that the cytokine response is minimally dependent on differentiation status although all major subets such as naïve, central memory, effector memory and effector cells are all detected. Antigen-specific activation increased the level and frequency of cells expressing immune checkpoints e.g., CTLA-4 and PD-1, and slightly reduced the frequency of co-stimulator expression, which correlates with elevated immunosuppressive cytokines like IL10 and TGFβ. All together, the activation states of these CAR-T cells are highly mixed with T_H1_, T_H2_, T_reg_, and GM-CSF-producing T-cell responses in the same single cells and largely independent of differentiation status. This work provided the first comprehensive portrait of CAR-T activation states, which may differ between patients but our work does teach the general rules regarding how different subtypes of CAR-T cells respond to antigen-specific challenge. It provides valuable information about the third-generation CD19-BB-28-3z CAR-T cells, which are being used in clinical trials in order to further improve therapeutic efficacy for non-responding patients. Our work provides new insights to the biology of CAR-T cell activation and a route to develop single-cell approaches for CAR-T infusion product quality assurance and to monitor the changes of CAR-T cells in patients post infusion in clinical trials.

## METHODS

### Cell Culture and Labeling

Human anti-CD19scFv-CD28-4-1BB-CD3ζ (Promab) T cells were cultured in complete X-Vivo 10 (Lonza) medium supplemented with IL-2 (10 ng/mL, Biolegend) at a concentration of 0.5 x 10^6^ cells/mL. CD4/CD8 subsets were identified using anti-CD4 FITC (Miltenyi Biotec) and anti-CD8 Alexa Fluor 647 (Cell Signaling). To summarize, 10×10^4^ CAR cells were pelleted and resuspended in 100 ul of PBS with the above antibody cocktail. The labeling reaction was incubated for 15 minutes at room temperate away from light. The cells were then rinsed twice in PBS and once in complete RMPI medium with 10% FBS (Zenbio) (vol/vol). Raji cells were labeled with Vybrant DiD (Invitrogen, V22887) following manufacturer instructions. Briefly, the Raji cells were resuspended at a density of 1 x 10^6^ cells/ ml in serum-free RPMI containing 5 ul/mL of Vybrant DiD solution and incubated for 20 min at 37 °C. The labeled cells were then washed three times in complete RMPI medium.

### CAR-T cell stimulation with target tumor cells

CAR-T and Raji cells were cultured as described above. Prior to loading CAR cells and Raji cells (50×10^3^ cells/mL concentration) were incubated for 6 hours in a round bottom well plate. Following incubation, Raji cells were separated from co-culture by positive selection using a modified protocol of B cell selection by MagCellect (R&D Systems). Briefly, CAR:Raji co-culture was centrifuged and the cell pellet was resuspended in serum free RPMI and labeled with anti-CD19 biotinylated antibodies. Following manufacturer’s protocol, MagCellect Streptavidin Ferrofluid was added to the solution and incubated for the recommended amount of time. The Streptavidin Ferrofluid beads bound to the anti-CD19 biotinylated Raji cells are collected on the side of the test tube by applying a magnet and then the CAR cells are pipetted out of the reaction. This procedure is repeated twice to make sure that most Raji cells are separated. The isolated CAR cells are then added to the microarray device for mRNA capture and transcriptome sequencing. Unstimulated CAR-T cells (50×10^3^ cells/mL concentration) were incubated for 6 hours alone and then ready for single-cell RNA-seq.

### Bulk LDH cytotoxicity assessment

Cytotoxicity was assessed using the Pierce LDH Cytotoxicity Assay kit (Thermo). The effector: target ratios surveyed are 1:1, 5:1, and 10:1. Target and effector cells were maintained in complete RPMI medium at a final concentration of 0.7×10^6^ cells/ml. LDH release was measured in the supernatant according to manufacturer’s instructions. Briefly, effector and target cells were incubated as described above for a six-hour period. Maximum LDH release was measured in target cells incubated with the provided 10X lysis buffer. Target cell cytotoxicity was calculated using the following formula: %Cytotoxicity = 100*[(Car^+^: Raji – Car^+^ alone – target alone)/(Maximum target lysis – target alone)].

### Population Sytox Green Cytotoxicity Assay

Effector CAR-T cells and labeled target cells were co-cultured in a 96 well U-bottom plate. Three effector: target concentrations were surveyed: 1:1, 5:1, and 10:1. Raji cell concentration was kept at 10,000 cells/ml while the concentrations of CAR^+^ cells were as follows 10,000 cells/ ml, 50,000 cells/ ml, and 1,000,000 cells/ml for each different ratio mentioned above. To distinguish cell death, SYTOX (green) (Invitrogen) was added to each well according to manufacturer’s instructions. Images were taken at 0, 4, 6, 10 hours using the Nikon Eclipse Ti microscope.

### Single CAR-T cell cytotoxicity assay in a microwell array

100 µm x 100 µm nanowell array was prepared by curing poly(dimethyylxiloxane) PDMS (Dow Corning) on a silicon wafer master with the etched array design. The nanowell array was cut to fit within the well of a 24 well plate and it was surface treated with oxygen plasma prior to cell loading to decrease PDMS hydrophobicity^36^. Labeled Raji (T) and CAR cells (E), as described above, were seeded at a density of 100,000 cells/ ml. Stochastic distribution of the cells within the array allowed for a range of E:T ratios. Images were acquired on a Nikon Eclipse Ti microscope fitted with an incubation chamber (37 °C, 5% CO_2_) using a 10x/0.3 objective. Automated multi-loci images were taken at 15 min intervals for a total duration of 10 hours. Images were processed using the built-in Nikon software. Paired t-tests were used to determine *P* values as described in the legend of Figure 2.

### Fabrication of antibody barcode array slides and single-cell microchamber array chip

A panel of 14 antibodies (IL-2, IL-4, IL-5, IL-6, IL-8, IL-10, IL12p70, IL-13, IL-17a, Granzyme-b, TNF-α, TNF-β, IFN-γ, GM-CSF, and FITC-BSA (alignment control) (R&D) was patterned onto a poly-l-lysine glass slide using an air-pressure driven flow through a PDMS mold with 20 microfluidic channels. The PDMS mold was fabricated in-house as previously described^17, 18^. The subnanoliter microchamber array for cell capture contains 14 columns of 220 wells and was fabricated into a PDMS slab. Prior to cell loading the microchamber PDMS slab was treated with oxygen plasma and then topped with RPMI medium.

### Single-cell cytokine secretion profiling using antibody barcode chips, data collection and analysis

Following labeling, CAR and Raji cells were incubated at a 1:1 ratio for 15 minutes in RPMI medium supplemented with SYTOX (green) to final concentration of 0.2 µM and the suspension was then gently pipetted on the microchamber array. The SCBC was assembled by overlaying the 14-plex antibody barcode glass slide and secured using our clamping system. The SCBC assay was imaged (Nikon Eclipse Ti) once at 0 hours and then following a 14-hour incubation (37 °C, 5% CO_2_) was imaged a final time. Cell number (CD4+ CAR, CD8+ CAR, and Raji cells), cell-to-cell contact, and Raji death was recorded for each microchamber from the phase-contract and fluorescent images. After the 14 hour incubation, the antibody barcode chip was developed using an ELISA sandwich immunoassay as previously described^18^. The developed slides were scanned using the GenePix 4200A microarray scanner (Axon).

### Single-cell cytokine profile data quantification and informatics analysis

The fluorescent signals were processed using the GenePix software and a custom Excel macro to determine the average fluorescent signal for each secreted cytokine. The threshold gate (background) used for cytokine intensity normalization was calculated by (average of raw mean fluorescence intensity of a given cytokine for the 0-cell wells) + 2× (standard deviation of raw mean fluorescence intensity of a given cytokine for the 0-cell wells)^37^. Secretors were defined as cells in the microchambers where the corresponding fluorescence cytokine signal intensity was higher than the threshold. We performed log transformation on the normalized cytokine intensity values (secretor raw fluorescence intensity for each cytokine – threshold gate for each cytokine) and used analysis and graphical tools applied single-cell analysis and graphical tools to visually represent such large-scale high-dimensional data such as PCA and viSNE^38^. All single-cell secretomic analysis was performed using our custom-built algorithms in Excel, R, and MATLAB.

### Population-level cytokine secretion measurement of CAR : Raji cell pairings

CAR-T cells were co-incubated with target cells, both at a density of 100,000 cells/ml, in triplicates in a u-bottom 96 well plate at a 1:1 E:T ratio. CAR-T and Raji alone controls were also included in the same well plate at the same concentration. After a 10 hour incubation, the cell suspension was pelleted at 200 g and the supernatant was removed and the population cytokine assay was performed as previously described.^17, 18^

### High-throughput scRNA-seq using a microwell array chip and data analysis

Our approach for massively parallel 3’ mRNA sequencing of single cells was based upon a closed microwell array system for co-isolation of single cells and DNA barcode chip. The chemistry was modified from the previous publication^15^. Briefly, the cell suspension prepared as above was pipetted onto the inlet of our the flow cell of our microchip, in which ~15,000 microwells (40 micrometers in diameter) covered the entire bottom to allow for trapping of single cells. The cells were withdrawn into the device using a negative pressue applied to the outlet. Once the microwell array was filled with the cell solution, the fluid flow was stopped and cells were allowed to settle by gravity into the wells. Excess cells were washed out by PBS and mRNA capture beads were then loaded similarly to the cells. Excess beads were also washed out by PBS and then lysis buffer was introduced into the device. Fluorinated oil (Fluorinert FC-40) was withdrawn into the device to seal the wells. Following lysis, the microfluidic device was incubated for an hour inside a humidity chamber to allow mRNA capture onto the beads. Reverse transcription, library construction, and sequencing were performed as previously discussed. Sequencing libraries were constructed according to the Nextera tagmentation (Nextera XT, Illumina) manufacturer’s protocol. The finished libraries were sequenced on the HiSeq 2500 sequences (Illumina). The transcripts were then aligned to the reference human transcriptome hg19 using STAR v2.5.2b. The resulting gene expression matrices were analyzed using the Seurat package in R studio, custom built algorithms in R, MATLAB, and Excel and GraphPad Prism was used for plotting.

## Acknowledgments

This research was supported by the Packard Fellowship for Science and Engineering (R.F.), National Science Foundation CAREER Award CBET-1351443 (R.F.), U54 CA193461 (R.F.), U54 CA209992 (Sub-Project ID: 7297 to R.F.), Yale Cancer Center Co-Pilot Grant (to R.F.). Sequencing was performed at the Yale Center for Genome Analysis (YCGA) facility. The molds for microfluidic devices were fabricated in the Yale School of Engineering and Applied Science cleanroom. We graciously thank Michael Power, Chris Tillinghast and James Agresta for their help with fabrication process.

